# Growth phase influences virulence in *Candidozyma auris* systemic infection models

**DOI:** 10.1101/2025.06.02.657403

**Authors:** Michael J. McFadden, Juliet A.E. Anku, Faith A. Davis, Catherine Luke, Andrea Obi, Teresa R. O’Meara

## Abstract

*Candidozyma auris* is a growing public health concern, capable of causing long-term contamination of healthcare settings, skin colonization, and life-threatening bloodstream infections. However, *C. auris* pathogenesis is not well understood, which is exacerbated by limitations and discrepancies in existing animal infection models. Further, the effects of *C. auris* growth phase on virulence have not been examined, despite growth phase being linked to virulence in many bacterial species. To address this question, and to develop an immunocompetent murine model of infection, we directly compared log and stationary phase *C. auris* systemic infection in immunocompetent C57BL/6J mice at high and low doses of infection. Systemic infection with high dose log phase *C. auris* results in rapid mortality between 2 hours and 1 day post infection, whereas stationary phase *C. auris* results in significantly extended survival. However, at low doses of infection, there was no difference in mortality kinetics between log and stationary phase cells. We observed that *C. auris* initially colonizes multiple organs but is rapidly cleared from the lungs and spleen, while kidney fungal burdens remain stable. Mice infected with high dose log phase *C. auris* had Fibrin-associated blood clotting in multiple organs and decreased serum Fibrinogen levels, suggesting that coagulation may drive rapid mortality. This was associated with increased β-glucan exposure and mannan abundance in log phase *C. auris*. These results will inform the development of a more standardized animal model of systemic *C. auris* infection, which can be used to reveal key aspects of *C. auris* pathogenesis.

**Importance:** Despite its growing medical importance, there is limited understanding of *Candidozyma auris* pathogenesis, due in part to limitations of existing laboratory models of infection. To develop a more complete understanding of factors that contribute to *C. auris* pathogenesis, it will be necessary to establish consistent parameters for animal models of infection. To address this need, we directly compared log and stationary growth phases on *C. auris* pathogenesis in immunocompetent C57BL/6J mice using a single virulent Clade I isolate. At a high dose of infection, host survival was dramatically different between log or stationary phase *C. auris*, suggesting that growth phase can affect *C. auris* pathogenesis. These differences correlated with increased exposure of pathogen-associated molecular patterns in the *C. auris* cell wall in log phase cells. These results will be instrumental in the future development of standardized animal models to study *C. auris* pathogenesis.

## Introduction

*Candidozyma auris* is an often multidrug resistant invasive species within healthcare settings that also can cause systemic infection with mortality rates typically ranging from 30-60% (1, 2). *C. auris* pathogenesis is not well understood due to its recent emergence and the existence of comorbidities in many infected patients, as well as an incomplete understanding of host-pathogen interactions and pathology within sites of *C. auris* infection (3–5). While animal models have been of great utility in understanding the relative contribution of specific fungal mutants to virulence (6–9), there is not currently a standard animal model for *C. auris* disseminated disease. The immunocompromised model of *C. auris* was initially developed as a physiologically relevant model of systemic infection, with a particular focus on its utility for testing of antifungal compounds (10) or specific types of immunocompromise (11). Immunocompetent models of *C. auris* infection have also been developed in ICR outbred mice (12), BALB/c (13), and C57BL/6J (3, 4) backgrounds, with varying kinetics and rates of mortality. Additionally, *C. auris*-specific factors, such as strain background are likely a major source of variability, as evidenced by our recent work showing significant differences in virulence between two closely related clade I isolates (6, 14).

Beyond strain differences, work in bacterial pathogenesis has demonstrated that growth phase can also influence virulence of a single strain. However, the effects of growth phase are species-specific and were determined empirically. For example, for *Legionella* and *Brucella*, entry into stationary phase is associated with increases in virulence factor expression (15, 16), but for *Salmonella* and *Streptococci*, it is the exponential growth phase that is associated with virulence (17, 18). Therefore, we sought to determine the effects of *C. auris* growth phase and dosage on host survival and pathology using a single strain of *C. auris* and a single genetically tractable and immunocompetent murine model of systemic infection.

At a high dose of systemic *C. auris* infection, we observed that log/exponential phase fungi cause rapid mortality compared to stationary phase fungi, which cause mortality over the course of several days. These differences in mortality may stem from rapid extensive blood clotting caused by log phase *C. auris*, as we observed decreased serum Fibrinogen levels and increased presence of blood clotting and fibrin deposition in multiple organs. *In vitro* exponentially growing *C. auris* cells had increased β-glucan exposure and mannan abundance, potentially promoting detection by host cells and triggering blood clotting and rapid mortality. However, differences in mortality observed at a high dose of infection were ablated at a low dose of infection, in which mice survived over the course of multiple weeks. We recovered *C. auris* from the lungs, spleen, and kidneys early after infection, but over time, only the kidneys maintained a substantial fungal burden. Together, these results suggest that growth phase can have dramatic effects on survival during systemic *C. auris* infection. Additionally, our work provides new insight into *C. auris* disease progression and will be helpful in establishing more standardized approaches to modeling *C. auris* infection in immunocompetent mouse models.

## Results

### Comparing the Effect of *Candidozyma auris* Growth Phase on Survival Kinetics and Fungal Burden in Organs

To assess whether growth phase affects virulence during systemic *C. auris* infection, we compared infection outcomes in an immunocompetent murine model using log or stationary phase *C. auris* cultures at previously published high and low doses of infection (5×10^7^ and 1×10^6^, respectively) using the virulent AR0382 (B11109 clade 1) isolate (Fig. 1A). Following intravenous infection, each group was monitored for several hours for the onset of disease symptoms. A majority of the cohort infected with a high dose of log phase *C. auris* rapidly declined in health, becoming moribund and showing labored breathing within 2 hours of infection, resulting in humane sacrifice. None of the mice in this group survived beyond the first day post-infection (Fig. 1B). In contrast, mice infected with the same high dose of stationary phase *C. auris* survived significantly longer than their log phase-infected counterparts, with onset of mortality starting at day 2 post-infection and full cohort mortality only occurring at day 6 post-infection (Fig. 1B). At the lower dose of infection, we did not observe any significance in mortality between the log and stationary phase infection cohorts (Fig. 1C). These data suggest that the growth phase of *C. auris* affects pathogenesis in murine infection models specifically at a high dose of infection.

**FIG 1:**
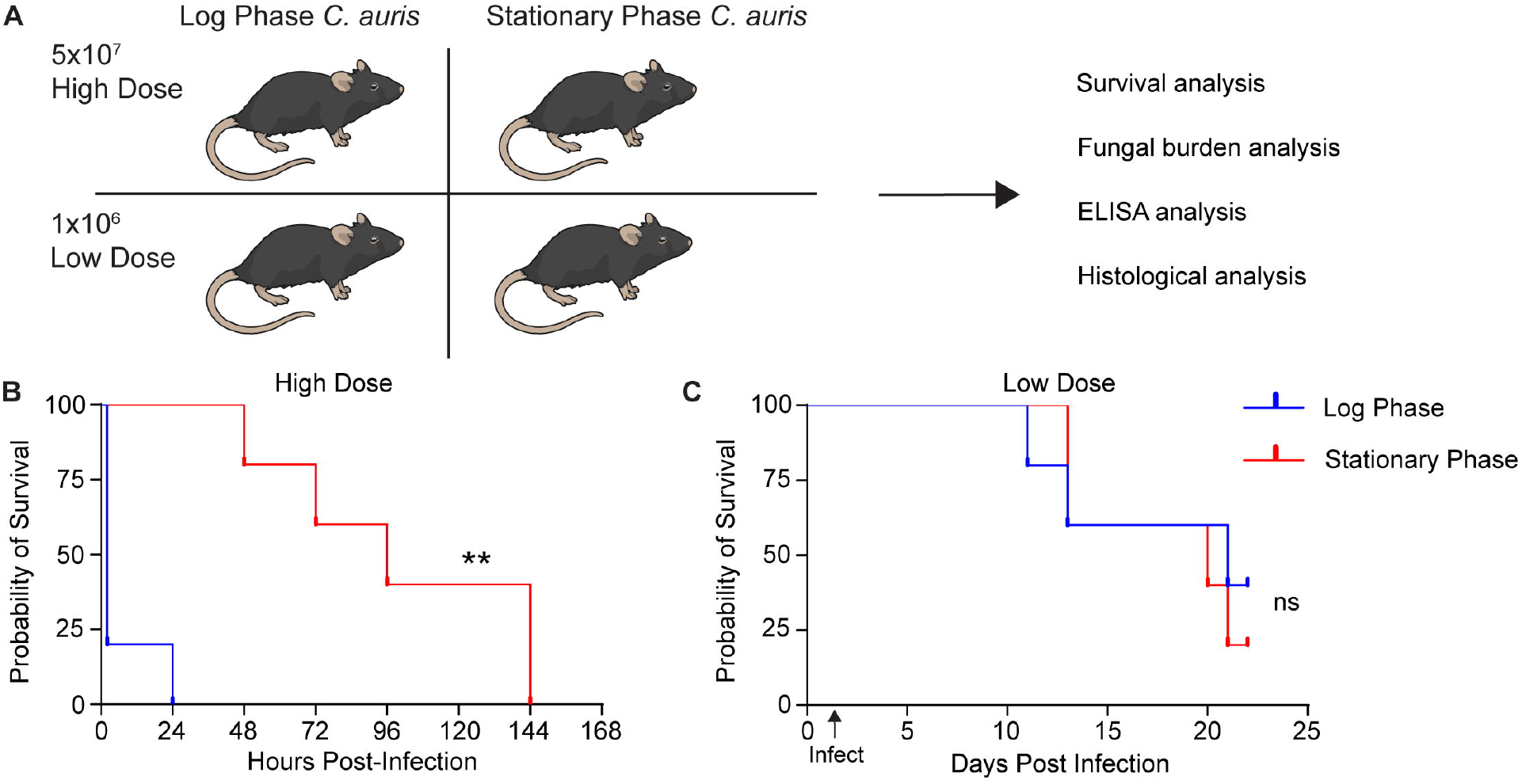

We measured the fungal burden in the lungs, spleen, and kidneys post-mortality in each cohort. In the high dose log phase *C. auris*-infected cohort, which had very rapid mortality, we observed a significantly higher fungal burden in the lungs than in the stationary phase cohort, while there was no difference in burden in the kidneys or spleen (Fig 2A-C). However, we hypothesized that the difference observed in lung fungal burden is likely a product of survival kinetics, rather than colonization, as the fungal burden in the lungs appeared to sharply decrease over time (Fig. 2D). To test this, we performed a matched comparison of *C. auris* burden in the lungs at 2 hours post-high dose infection for both log and stationary phase and observed similar burden between both cohorts (Fig. 2G). Additionally, in low dose cohorts, which survived significantly longer than high dose cohorts, we recovered very few fungal colonies from the lungs from both cohorts, indicating that *C. auris* is effectively cleared from the lungs over time (Fig. 2H).

**FIG 2:**
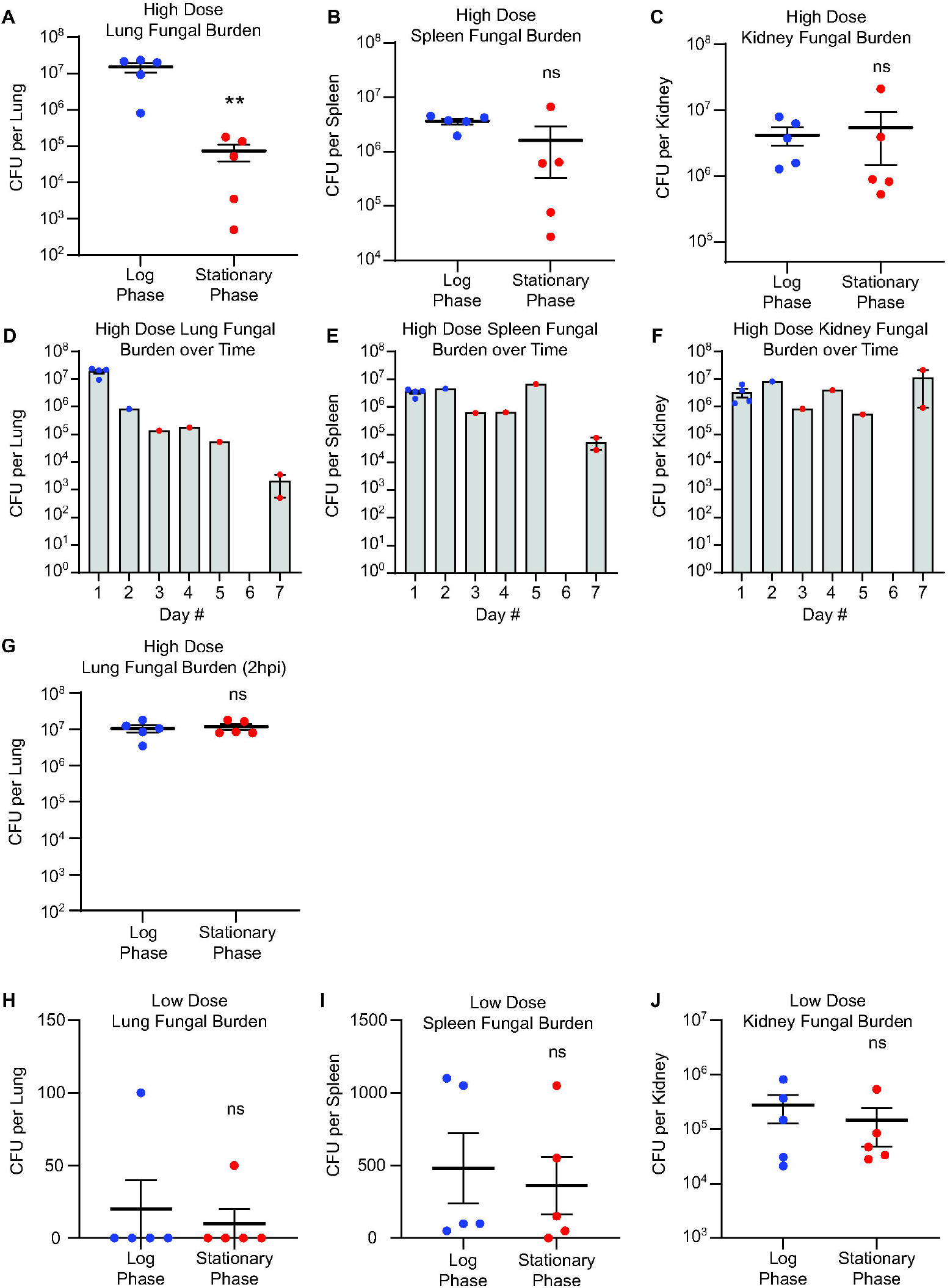

Similarly, fungal burden in the spleen showed a decreasing trend at day 7 in the high dose cohort (Fig. 2E) and was generally low at the time of mortality in the low dose cohorts (Fig. 2I), consistent with clearance from these organs over time. In contrast, fungal burden in the kidneys remained consistent over time in both the high dose (Fig. 2F) and low dose cohorts (Fig. 2J). Together, these data suggest that *C. auris* disseminates to multiple organs after systemic infection but is cleared from the lungs and spleen, while fungal burden remains steady over time in the kidneys.

### High Dose Systemic Infection with Log Phase *C. auris* Causes Rapid Blood Clotting

We next sought to further understand the differences in mortality observed between log and stationary phase *C. auris* during systemic infection at a high dose. We noted animals that rapidly succumbed within the log phase high dose cohort exhibited blood clotting while performing cardiac puncture. Coagulation is associated with septic shock, which has a high mortality rate in the absence of intensive care and is associated with increased mortality in candidemia patients (1, 19). ELISA data from the 2 hour post-infection mice showed decreased Fibrinogen levels in the plasma of animals infected with either high dose log or stationary phase *C. auris*, compared to control mice (Fig. 3A), consistent with Fibrinogen being converted to insoluble Fibrin to form clots (20). However, animals infected with stationary phase cells recovered from this initial response and showed higher Fibrinogen levels at the time of mortality (Fig. 3B). Additionally, using Martius Scarlet Blue (MSB) staining in which Fibrin stains red, erythrocytes stain yellow, and muscle and collagen stain blue, we observed increased Fibrin deposition only in the log phase-infected animals, suggesting that the coagulation cascade proceeded in this cohort to trigger systemic thrombosis (Fig. 3C-F; Fig. S1). Quantification of Fibrin staining revealed that only log phase-infected kidneys showed significantly higher Fibrin levels relative to erythrocyte abundance in MSB-stained kidney sections (Fig. 3G). The mock and stationary-phase kidneys at both time points were not statistically different (Fig. 3G). Blood clots could also be observed in MSB-stained lungs from the log phase infected cohort at the time of mortality (Fig. S2), consistent with systemic thrombosis in log phase-infected animals, rather than in a specific organ.

**FIG 3:**
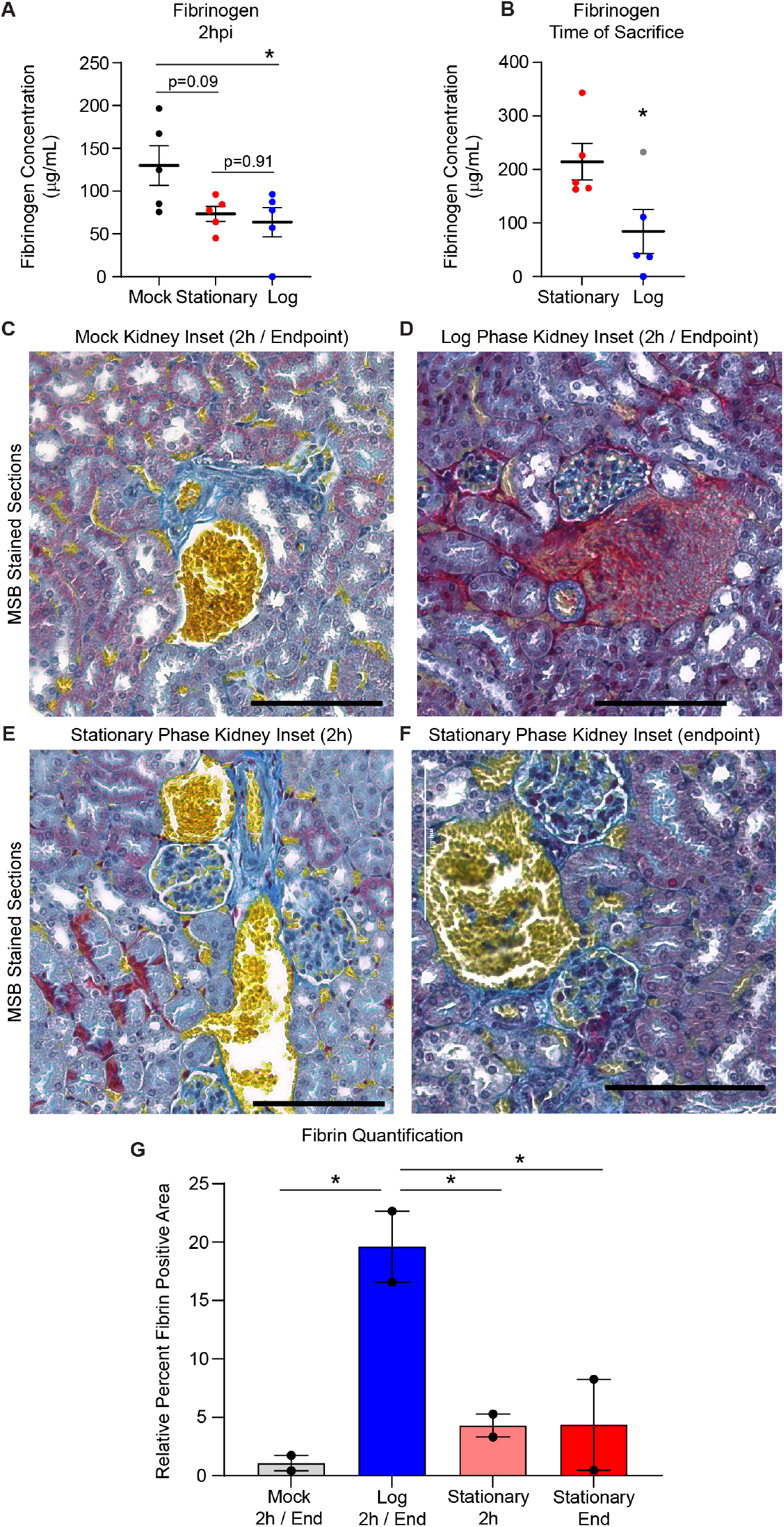

To determine whether a proinflammatory response drives mortality and coagulation in the high dose log phase-infected cohort, we measured the levels of proinflammatory cytokines in plasma samples from the high dose infection cohorts at a synchronized time of 2 hours post-infection (hpi) and post-mortality. We did not observe consistent signs of systemic inflammation in either log or stationary phase high dose infection cohorts at 2 hpi, as TNF levels were only elevated in stationary phase infected mice (Fig. 4A), IL-6 levels were only elevated in log phase infected mice (Fig. 4B), and IL-1β levels were unchanged compared to mock-infected control mice (Fig. 4C). Proinflammatory cytokine levels were also similar at the time of mortality between log and stationary phase high dose cohorts (Fig. 4D-F), again consistent with blood clotting, rather than a cytokine storm, driving rapid mortality after systemic infection with log phase *C. auris*. Hematoxylin and eosin (H&E)-stained lungs showed blood clots specifically in log phase-infected lungs, but not dramatic immune cell recruitment (Fig. 4G), consistent with these findings. Together, these data suggest that rapid and aberrant systemic coagulation causes mortality after infection with high doses of log phase *C. auris*.

**FIG 4:**
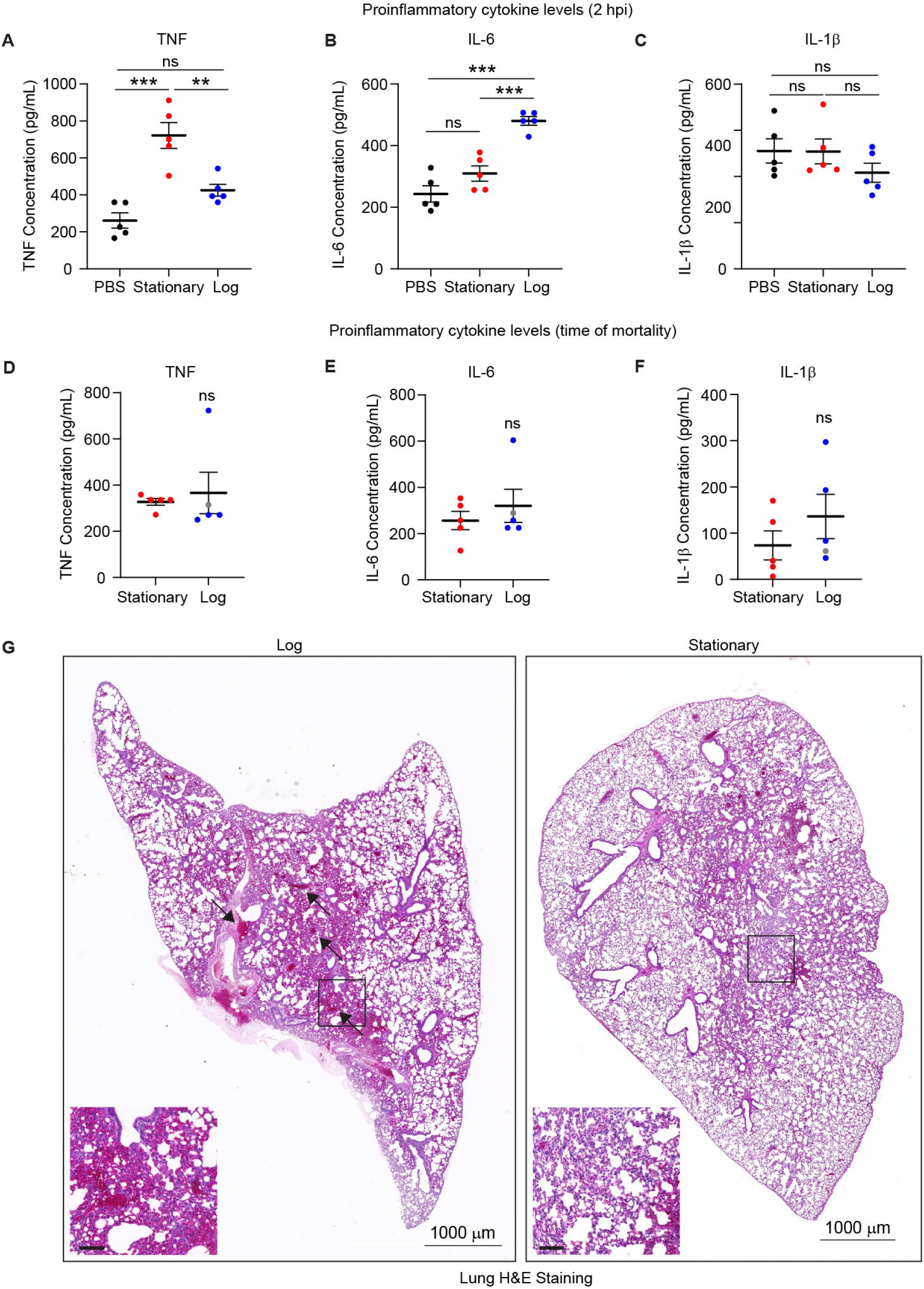

We then hypothesized that *C. auris* growth phase would affect the composition of the cell wall, triggering differential recognition and leading to systemic coagulation. Therefore, we measured the major cell wall components and fungal pathogen-associated molecular patterns (PAMPs) chitin, mannan, and exposed β-glucan in log or stationary phase *C. auris*. Flow cytometry analysis revealed that log phase *C. auris* showed higher β-glucan exposure than stationary phase (Fig. 5A, D). Interestingly, there were two distinct populations of high and low mannan cells in the log phase cells, compared to stationary phase cells, which showed only a single peak with intermediate intensity (Fig. 5B, E). In contrast, chitin levels were similar between log and stationary phase cells (Fig. 5C, F). Similar results were observed through confocal microscopy, which revealed increased β-glucan exposure on log phase *C. auris* cells and a subset of cells with increased mannan staining enriched on the cell periphery. Chitin levels were heterogeneous between cells but overall levels did not differ between stationary and log phase cells (Fig. 5G). These results reveal differences in the composition of the cell wall and exposure of pathogen associated molecular patterns in log phase *C. auris* cells, which may drive the immunopathological response and mortality observed during systemic infection at a high dose.

**FIG 5:**
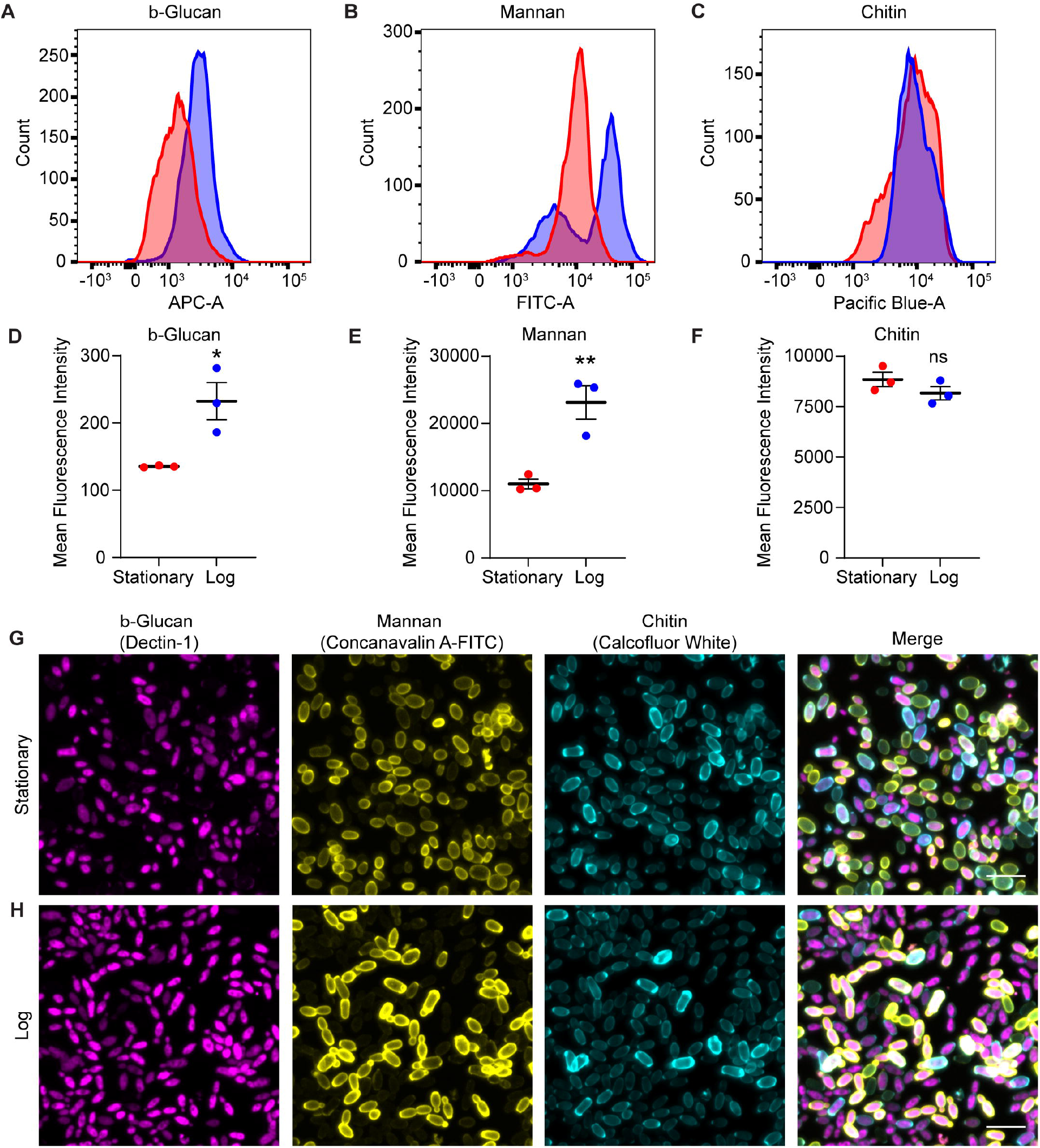

## Discussion

Here, we show that the growth phase of *C. auris* influences its virulence in a murine model during bloodstream infection at a high dose of infection. Notably, the inoculum used in our high dose cohort has been commonly used in previous reports for *in vivo* models of *C. auris* pathogenesis and survival analyses (6, 8, 9). Therefore, growth phase will be an important parameter to consider for future research on *C. auris* pathogenesis and a crucial consideration in establishing animal models. Growth phase is often associated with virulence programs in bacteria (15–18), and our results suggest that log phase growth in *C. auris* is associated with increased exposure of mannan and β-glucan, which are major fungal PAMPs. These cell wall changes correlate with initiation of blood clotting during bloodstream infection in a murine model of infection, leading to rapid mortality, potentially caused by septic shock, systemic thrombosis, and disseminated intravascular coagulation. Indeed, we found decreased Fibrinogen levels in plasma samples from mice infected by log phase *C. auris* at a high dose, as well as increased Fibrin deposition in the kidneys and lungs. These results suggest a systemic thrombosis response specifically to log phase *C. auris* at a high dose, which we propose causes rapid mortality.

Understanding the cause of the observed blood clotting response will be an important future direction of this work. While we observed increased exposure of mannan and β-glucan, additional work will be needed to establish these cell wall changes as the fungal determinant of coagulation in murine models. Mannan binding lectin-associated serine proteases can trigger blood clotting (21), revealing one possible mechanism by which the high mannan log phase *C. auris* may trigger rapid mortality at a high dose. Whether these cell wall alterations are conserved in other strains of *C. auris*, which can differ in cell wall composition and virulence (22), and whether additional strains also cause thrombosis in log phase growth are also significant outstanding questions. Additionally, elucidating the host cell populations responsible for triggering coagulation will be of interest, as resident or circulating innate immune cells, such as monocytes, macrophages, and neutrophils can detect fungal PAMPs and produce coagulation factors (23–25). However, vascular endothelial cells have also been shown to express PRRs that can recognize fungal cell wall components (26, 27), and thus may play an important role in thrombosis during *C. auris* bloodstream infection.

By comparing fungal burden in the organs of mice post-mortem, we found that *C. auris* burden is high in the lungs early after infection but is rapidly cleared, whereas *C. auris* establishes a more stable burden in the kidneys. The kidneys are also considered the primary target organ of *C. albicans* (28, 29), although it is not immediately clear why the kidneys are a more permissive environment for *C. auris* colonization than the lungs. However, future work to reveal how the lungs clear *C. auris* may lead to new understanding of effective organ-specific immune responses against *C. auris* and perhaps fungal pathogens more broadly.

Full understanding of the clinical implications of this report is hindered by a lack of knowledge of what growth state is common for *C. auris* when initiating bloodstream infections in humans. However, septic shock may occur in more than 20% of patients with *C. auris* candidemia and increases mortality, and thrombosis has also been reported during *C. auris* candidemia (1, 30, 31). Therefore, understanding the molecular mechanisms by which *C. auris* infection causes thrombosis during bloodstream infection may inform therapeutic strategies to treat these high-mortality infections. Together this work establishes virulence differences between stationary and log phase *C. auris* at high doses and indicates that the development of a standardized murine infection system, including controlling for fungal growth phase, will be important for future studies examining *C. auris* pathogenesis.

### Limitations of the Study

This report has several limitations that will be important to address in future studies. While this work focuses on virulence differences between log and stationary phase growth *C. auris* cells during bloodstream infection at a high dose, these differences were not readily seen at a lower dose of infection. Thus, it is unclear how these results translate to human infection, although thrombosis has been reported in patients with *C. auris* candidemia. Additionally, this study focuses primarily on a single strain of *C. auris* and the widely used C57BL6/J mouse model, thus additional work is needed to determine whether the observed phenotypes are generalizable across *C. auris* strains and clades and with different animal models. While we report increased abundance of mannan and exposure of β-glucan in the *C. auris* cell wall during log phase growth, which correlates with blood clotting observed during systemic infection, we have not definitively shown whether these cell wall changes are responsible for triggering coagulation. While our results support that systemic thrombosis is the likely cause of rapid mortality in high dose log phase *C. auris*-infected mice, we have not extensively profiled tissue-specific inflammation, thus we cannot definitively rule out a proinflammatory response within specific organs. Finally, the host cells that initiate the coagulation response to log phase *C. auris* infection are not identified in this work.

## Methods

### *Candidozyma auris* growth conditions

*C. auris* strain CDC-AR0382 (B11109) was grown in YPD liquid media (1% yeast extract, 2% peptone, 2% dextrose) with constant agitation. After 16 hours, stationary phase cultures were sub-cultured by diluting to OD600 of 0.2 in fresh YPD and grown at 30°C for 4 hours with constant agitation to establish log phase growth.

### Survival analysis post-systemic infection

*C. auris* from log phase or stationary phase cultures were pelleted by centrifugation (5000 rpm for 5min), washed once with sterile PBS, and resuspended in sterile PBS to desired doses for infection. Immunocompetent 8-week old female C57BL/6J mice were infected intravenously with *C. auris* from log phase or stationary phase growth at high dose (5×10^7^) or low dose (1×10^6^) via retro-orbital injection in 100 μL volume. Immediately following infection, mice were monitored for onset of disease symptoms for several hours initially, then daily over the course of 21 days. Mice were sacrificed at a humane endpoint defined as loss of 20 percent of initial bodyweight, or when severe disease symptoms were observed, such as unresponsiveness and labored breathing, or severe neurological symptoms.

### Analysis of fungal burden in organs post-mortality

After sacrifice, organs samples were harvested to measure fungal burden. The right lung, right kidney, and spleen were harvested by dissection and homogenized by bead beating with sterile ⅛ inch ball bearings (Grainger 4RJL3) for 10 seconds. Serial dilutions were performed and plated on YPD agar with ampicillin (100 mg/mL) and gentamicin (50 mg/mL). Fungal colonies were grown for 2 days at 30°C and counted, and fungal burdens per organ were calculated.

### Histological analysis and Martius Scarlet Blue staining for Fibrin analysis and quantification and in organs

After sacrifice, organs samples were harvested to perform histological analysis. The left lung and left kidney were harvested by dissection and fixed in 10 percent formalin for 24 hours, then plunged in 70 percent ethanol prior to sectioning and embedding on slides by the University of Michigan Orthopaedic Research Laboratories Histology Core. For histological analysis, hematoxylin and eosin (H&E) staining was performed by the University of Michigan Orthopaedic Research Laboratories Histology Core. Full organ images were captured at 10x magnification with color brightfield imaging using a BioTek Lionheart FX Automated Microscope. For Martius Scarlet Blue (MSB) staining, slides were deparaffinized through xylene and gradient ethanol (EtOH), and rehydrated to distilled water (diH2O). Postfix was performed in Bouin Fixative (Newcomer Supply, Inc., Middletown, WI) at room temperature overnight (~16 hours), followed by a 15 minute wash in diH2O. MSB staining was then performed, as previously described (32). Images of full organs were captured at 20x magnification with color brightfield imaging using a BioTek Lionheart FX Automated Microscope. For analysis of relative Fibrin levels, 6 representative erythrocyte-rich regions were blindly chosen from each organ and a pixel classifier was trained in QuPath v0.6.0 (33), using the wand tool to segment Fibrin-positive pixels, erythrocyte-positive pixels, and kidney tissue to ignore, in order to quantify Fibrin levels relative to erythrocytes within each image. The pixel classifier was then used for each image to quantify the Fibrin-positive pixel area, relative to erythrocyte-positive pixel area.

### Histological analysis in organs post-mortality

After sacrifice, organs samples were harvested to perform histological analysis. The left lung and left kidney were harvested by dissection and fixed in 10 percent formalin for 24 hours, then plunged in 70 percent ethanol prior to sectioning and staining (H&E and PAS) by the University of Michigan Orthopaedic Research Laboratories Histology Core. Slides were imaged using a BioTek Lionheart FX automated microscope.

### ELISA

After sacrifice, serum was collected by cardiac puncture, followed by isolation of serum using centrifugation (8000g for 5 minutes) of lithium heparin serum collection tubes (Kent Scientific KMIC-LIHEP). Serum samples were submitted to the University of Michigan Cancer Center Immunology Core for quantification of Fibrinogen, TNF, IL-6, and IL-1β by ELISA.

### Analysis of cell wall content

Log and stationary phase *C. auris* cells were pelleted by centrifugation (3000 x g for 5 minutes), washed once in PBS, and fixed in 4% paraformaldehyde for 15 minutes. Following fixation, *C. auris* cells were pelleted by centrifugation (3000 x g for 5 minutes), washed twice in PBS, and then stained for cell wall contents, followed by flow cytometry analysis or confocal microscopy. To quantify mannan content, cells were stained with 5 μg/mL of FITC-Concanavalin A (MilliporeSigma, C7642) for 30 minutes. To quantify exposed β-1,3-glucan, cells were blocked with 3% bovine serum albumin & 5% normal goat serum (Invitrogen, 10000C) for 30 minutes. After blocking, cells were stained with 15 μg/mL of hDectin-1a (InvivoGen, fc-hec1a-2) for 1 hour. Cells were washed twice with PBS before secondary staining with 4 mg/mL of goat raised anti-human IgG antibody conjugated with Alexa Fluor 647 (Invitrogen A-21445) for 30 minutes. To quantify chitin content, cells were stained with 0.1 g/L of Calcofluor White (MilliporeSigma, 18909-100ML-F) for 10 minutes. Following staining, cells were washed with 500μL PBS three times and resuspended in 500μL PBS. Samples were analyzed on a LSRFortessa Flow Cytometer (BD Bioscience, NJ, USA) using BD FACSDiva Software. 10,000 events were recorded for each sample. FlowJo software was used to determine mean fluorescence intensity. For confocal microscopy, images were captured on a Yokogawa CellVoyager CQ1 automated confocal microscope at 40x magnification, and maximum intensity projections were generated from confocal Z-stacks of 10 µm. Representative images were processed using NIH Fiji/ImageJ (34).

## Acknowledgements

We thank Joel Whitfield of the University of Michigan Cancer Center Immunology Core and Emma Snyder-White of the University of Michigan Orthopaedic Research Laboratories Histology Core (NIH P30 AR069620) for guidance and technical support for ELISA and histological analysis, and members of the O’Meara lab for discussion. This work was supported by National Institutes of Health grants U19AI181767 and the Burroughs Wellcome Fund Investigators in Pathogenesis award 1173374 to TRO.

