## Supplement for "Growth phase influences virulence in *Candidozyma auris* systemic infection models"

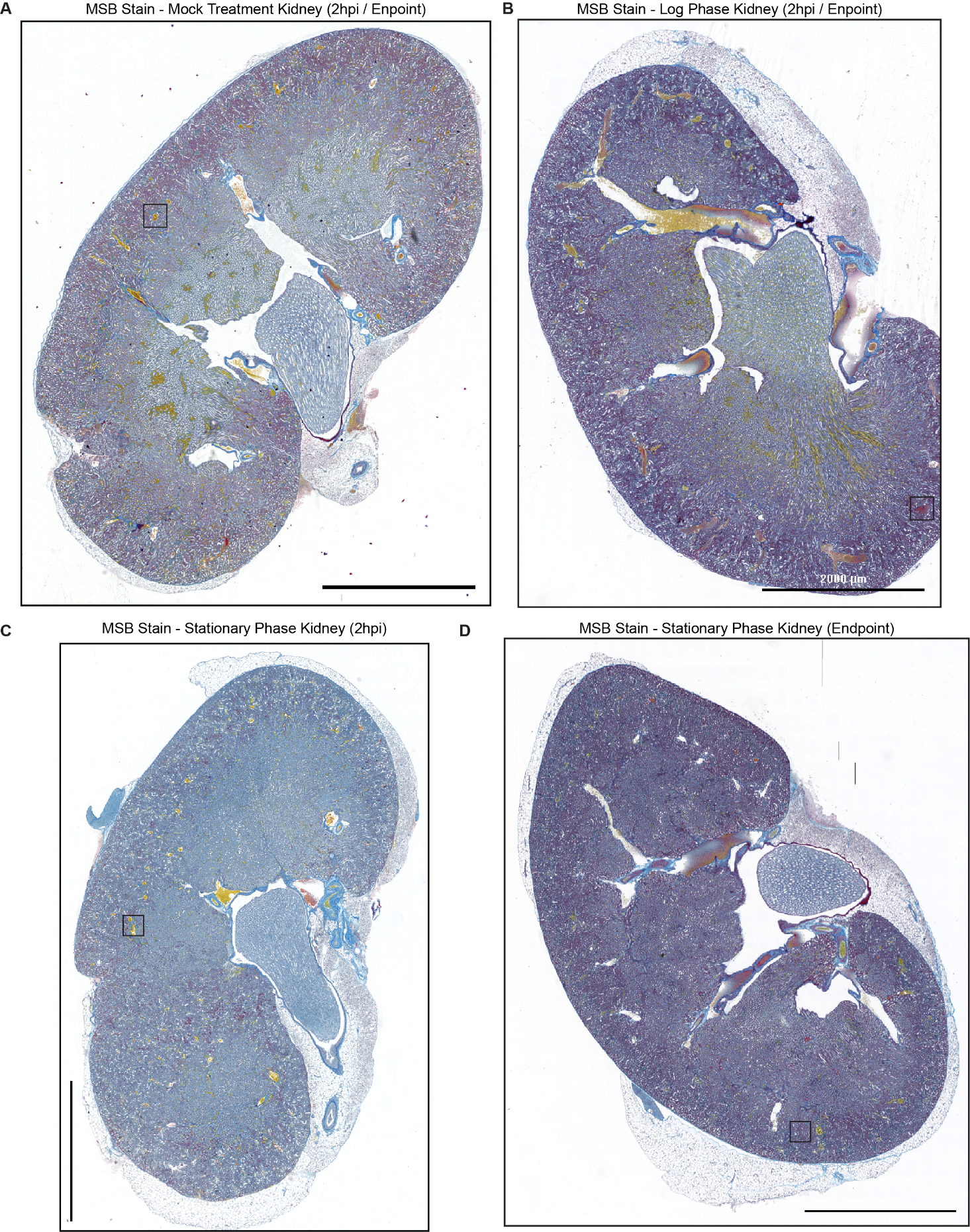

**Supplemental Figure 1: Related to Figure 3. (A-D)** Full-size images of Martius Scarlet Blue staining of kidney sections from mice after mock treatment (A), or high-dose infection with *C. auris* in log phase growth (B), or stationary phase growth at a matched timepoint of 2 hpi (C), or post-mortality (D) (scale bars 2000 μm). Images are representative of 2 animals per cohort. Box indicates region expanded in Fig 3.

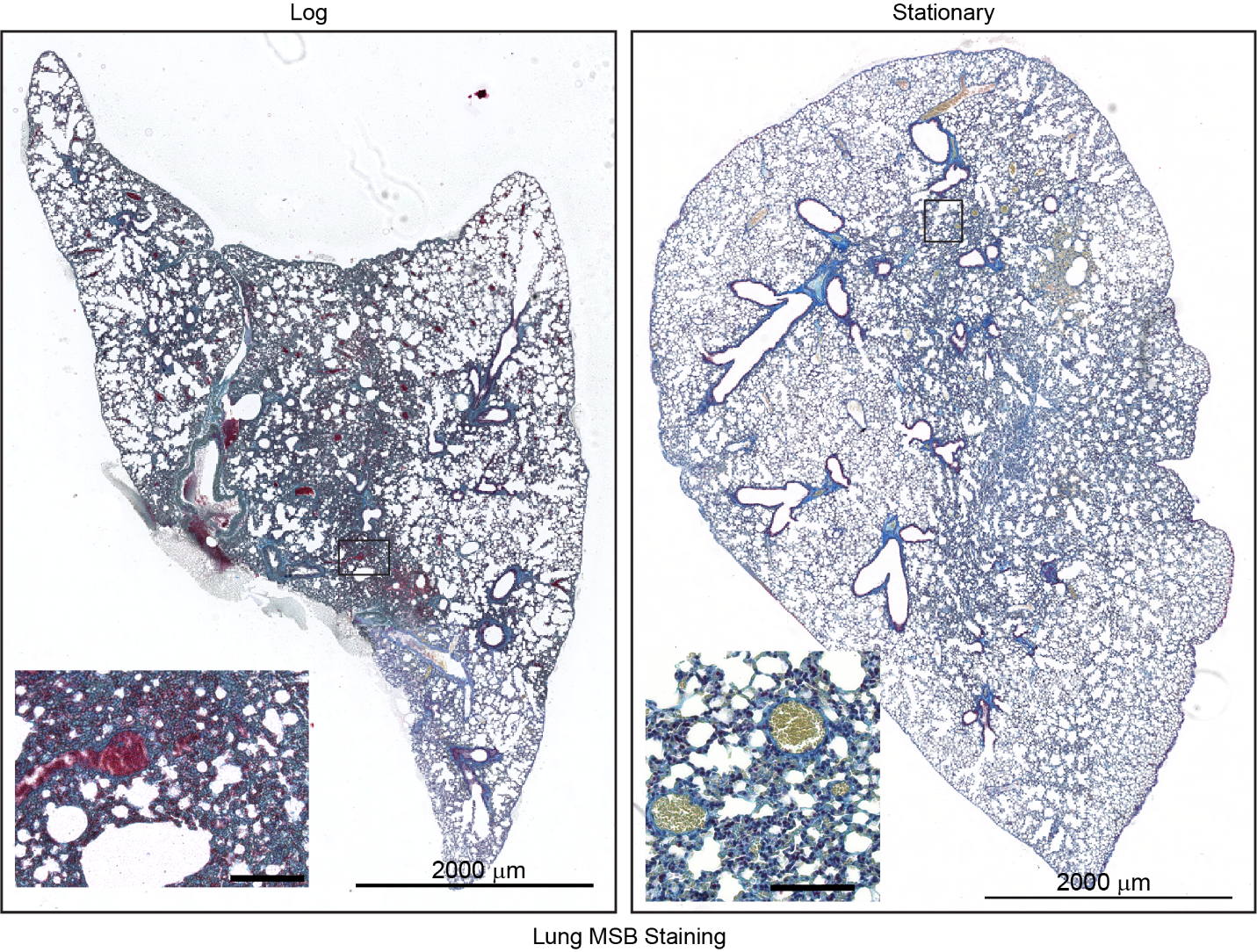

**Supplemental Figure 2:** **Related to Figure 3.** Representative MSB staining of lung sections from mice in high dose cohorts infected with log phase (left) or stationary phase (right) *C. auris* at time of mortality. Boxes show a magnified inset, showing Fibrin-positive erythrocyte-rich regions in log phase *C. auris*-infected lungs or Fibrin-negative erythrocyte-rich regions in stationary phase *C. auris*-infected lungs (scale bar 100 μm). Images are representative of 2 animals per cohort.
